# MitoROS due to loss of *Slc4a11* in corneal endothelial cells induces ER stress, lysosomal dysfunction and impairs autophagy

**DOI:** 10.1101/2020.08.27.250977

**Authors:** Rajalekshmy Shyam, Diego G. Ogando, Moonjung Choi, Joseph A. Bonanno

## Abstract

Recent studies from *Slc4a11* KO mice have identified mitochondrial dysfunction as a major contributor toward oxidative stress and cell death in Congenital Hereditary Endothelial Dystrophy. Here we asked if this stress activated autophagy in the *Slc4a11* KO cell line and in KO mouse endothelial tissue. Early indicators of autophagy, phospho-mTOR and LC3-II indicated activation, however P62 was elevated suggesting an impairment of autophagy flux. The activity and the number of lysosomes, the organelle responsible for the final degradation of autophagy substrates, were found to be reduced in the KO. In addition, the expression of the master regulator of lysosomal function and biogenesis, TFEB, was significantly reduced in the KO corneal endothelia. Also, we observed increased Unfolded Protein Response, as well as elevated expression of ER stress markers, BIP and CHOP. To test if lysosomal and ER stress stems from elevated mitochondrial ROS, we treated *Slc4a11* KO corneal endothelial cells with the mitochondrial ROS quencher, MitoQ. MitoQ restored lysosomal enzymes as well as TFEB, reduced ER stress, and increased autophagy flux. MitoQ injections of *Slc4a11 KO* mice decreased corneal edema, the major phenotype associated with CHED. We conclude that mitochondrial ROS causes ER stress and lysosomal dysfunction with impairment of autophagy in *Slc4a11* KO corneal endothelium. Our study is the first to identify the presence as well as cause of lysosomal dysfunction and ER stress in an animal model of CHED, and to characterize inter-organelle relationship in a corneal cell type.

## Introduction

The corneal endothelium is a monolayer on the posterior corneal surface facing the anterior chamber of the eye. These non-regenerating cells maintain corneal hydration and transparency through primary and secondary active transport processes ^1,2^. The energy needed for this process is derived from a high density of mitochondria. Corneal endothelial diseases, such as Fuchs Corneal Endothelial Dystrophy (FCED) and Congenital Hereditary Endothelial Dystrophy (CHED), are associated with dysfunctional mitochondria, increased reactive oxygen species (ROS), alterations in cell morphology, function, and eventually cell death ^3–6^. Corneal edema is the main clinical manifestation of FCED and CHED ^7,8^, and corneal transplantation is the only available treatment option. While CHED is a rare recessive disorder, FCED affects nearly 4% of people over the age of 40 and accounts for the majority of corneal transplantations across the world ^7–9^.

Corneal endothelial cells highly express Slc4a11, an electrogenic NH_3_ ^+^/H^+^ transporter, located in the basolateral membrane ^10^ as well as in the inner mitochondrial membrane ^5^. Homozygous recessive mutations in this gene are linked to early-onset CHED ^11^, whereas heterozygous mutations are implicated in later life FCED ^12^. A mouse model in which *Slc4a11* is knocked out emulates the disease progression of CHED along with increased oxidative stress in the corneal endothelium ^13,14^, a well-known characteristic of corneal endothelial dystrophies ^5,6,8^. Recent work from our lab demonstrated that Slc4a11 is an ammonia sensitive mitochondrial uncoupler. Absence of Slc4a11 causes glutamine-induced mitochondrial hyperpolarization, excess ROS production, deficient ATP production, and eventual apoptosis ^5^. Mitochondrial ROS can lead to increased autophagy which in certain cell types help in survival ^15^. Whether autophagy is activated by Slc4a11 loss of function, and if it enhances cell survival in the corneal endothelium are unknown.

In the current project we examined the autophagy process in corneal endothelial cells lacking Slc4a11 and found that while autophagy is activated, it is aberrant. We found that autophagy flux is disrupted in both the *Slc4a11* KO corneal endothelial cell line and mouse endothelial tissue, primarily due to lysosomal dysfunction, with concomitant ER stress and mitochondrial ROS. Quenching mitochondrial ROS with MitoQ rescued ER stress, lysosomal function, and autophagy flux. Moreover, MitoQ slowed the development of corneal edema in the *Slc4a11* KO mouse. These findings provide further evidence that mitochondrial Slc4a11 and the prevention of excess ROS production has a significant role in corneal endothelial dystrophy.

## Results

### Impaired autophagy flux in Slc4a11 KO corneal endothelial cells and tissue

Using immortalized mouse corneal endothelial cells (MCEC) established from *Slc4a11* wild-type (WT) and KO mice ^16^, we first asked if autophagy is activated. Increased autophagy is often triggered by a decrease in phosphorylated Membrane Target of Rapamycin (mTOR). This stimulates phagophore formation which in turn matures into an autophagosome and eventually fuses with lysosomes resulting in the degradation of its components as summarized in Figure 1A. Figure 1B and C show that the p-mTOR/mTOR ratio was decreased in *Slc4a11* KO cells. Similarly, the ratio of Microtubule associate protein 1A/1B Light chain 3 (LC3II/I) was increased (FIG 1B and C) suggesting that the initial steps of autophagy are activated in KO cells relative to WT. P62, a marker of autophagy flux that is part of the phagosome and typically low during activated autophagy, however was elevated in KO cells (FIG 1B and C). In cells with normal autophagy flux, treatment with Bafilomycin A1, which blocks vacuolar H^+^-ATPase activity, alkalinizes lysosomes and increases the accumulation of autophagy related proteins (FIG 1A). In WT cells, Bafilomycin increased the levels of Autophagy related 5 protein (Atg5), P62 and LC3II/I ratio (FIG 1B and C), as expected. However, in KO cells, this treatment did not elevate the expression of these proteins (FIG 1B and C). This result suggests a compromised autophagy flux in *Slc4a11* KO MCEC.

**Figure 1.**
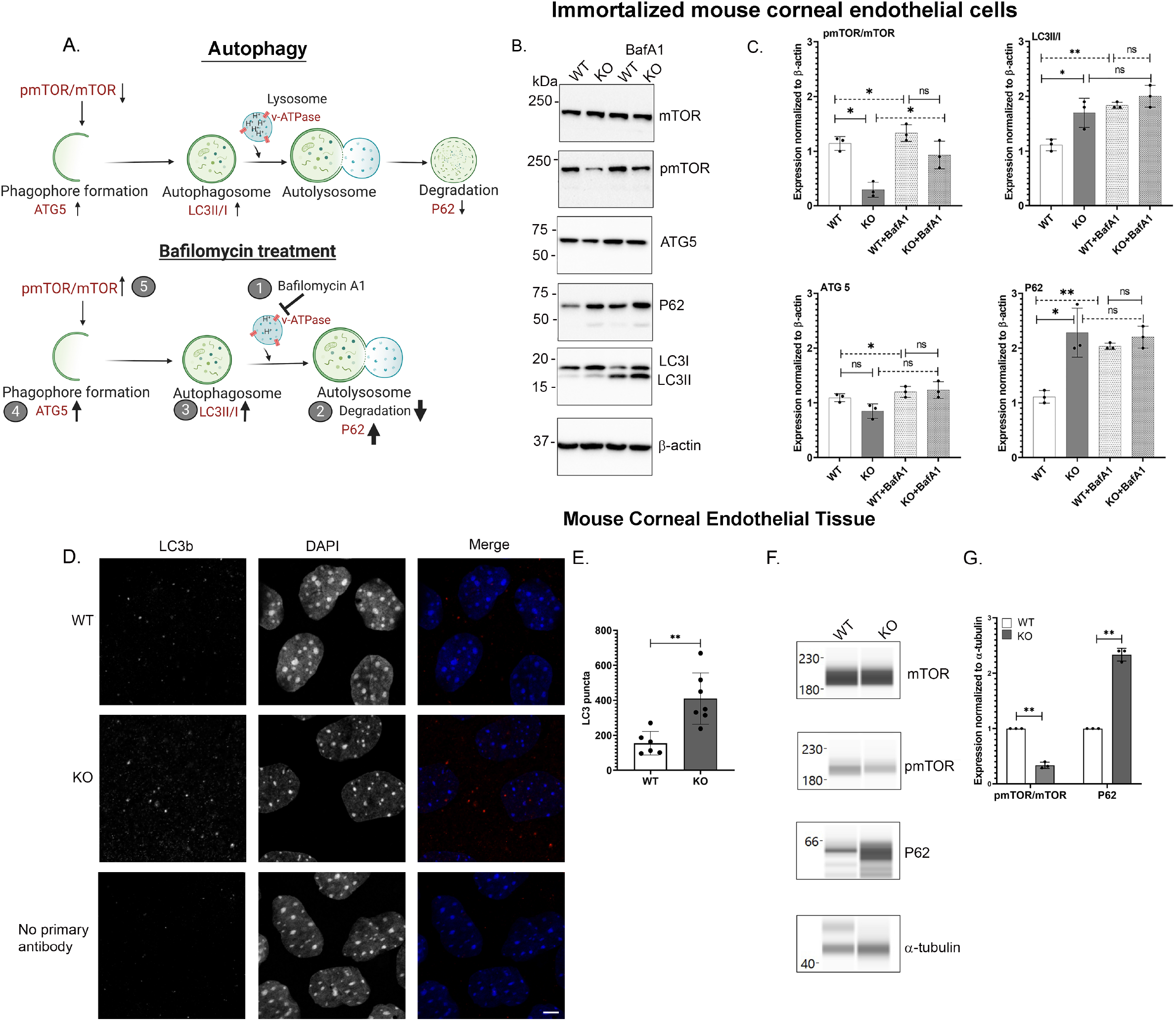
Autophagy flux is negatively affected in Slc4a11 KO corneal endothelial cells. **Panel A** In basal autophagy, autophagosome formation occurs followed by fusion with lysosomes and the degradation of substrates. In bafilomycin treated cells (1), the lysosomal proton pump (v-ATPase) which is responsible for its acidic environment is inactivated. This leads to the accumulation of undegraded autophagy substrates in the cell (2), which further increases the levels of proteins involved in autophagosome (3), and phagophore formation (4), which eventually increases pmTOR/mTOR ratio (5). **Panel A** Western Blots of *Slc4a11* WT and KO MCEC (treated with or without 50nM BafA1) showing the expressions of p-mTOR (Ser 2448), mTOR, Atg5, LC3II and P62. **Panel A** – Quantification of Western Blot data in Panel B. *p<0.01, **p<0.001, ns-not significant. **Panel A** Corneal endothelial tissue from *Slc4a11* WT and KO were stained with LC3b antibody (Red) and DAPI (blue). No primary antibody control shows minimal background staining. Scale bar – 5μm **Panel A** Quantification of LC3 puncta from Panel D. **p<0.001 **Panel A** WES protein analysis of corneal endothelial tissues from *Slc4a11* WT and KO animals showing the expressions of mTOR, pmTOR (Ser 2448) and P62. **Panel A** Quantification of WES analysis from Panel F. *p<0.01, **p<0.001

Next, we transfected the MCEC WT and KO cells with pMRX-IP-GFP-LC3-RFP-LC3ΔG, a fluorescent probe of autophagy flux ^17^. Here, the c-terminus of LC3 is cleaved by Atg4, an autophagy initiation factor. This results in the production of GFP-LC3 and RFP-LC3ΔG. GFP-LC3 is degraded in the lysosomes upon autophagy, whereas RFP-LC3ΔG remains cytosolic (FIG S1A). Using this method, autophagy flux can be assessed by comparing the GFP/RFP ratio. In the WT cells (FIG S1B and C), the GFP/RFP ratio was two-fold lower than that of the KO cells. Consistent with the Western Blot results, Bafilomycin caused an increased GFP/RFP ratio in the WT cells (FIG S1B and C), whereas in the KO cells the ratio did not change. Our data suggest that while there is autophagy induction in KO cells, autophagy flux is impaired.

We then asked if this aberrant autophagy is also present in the *Slc4a11* KO mouse corneal endothelium. In corneal endothelia isolated from 10-week-old KO animals, a two-fold increase in LC3 puncta numbers (FIG 1D and E) was observed when compared to age matched WT controls. In addition, Wes protein analyses revealed increased expression of P62 and decreased levels of p-mTOR/mTOR ratio in the KO animal tissue (FIG 1F and G). These results parallel the *in vitro* findings confirming that autophagy is activated, but aberrant in *Slc4a11* KO corneal endothelium *in vivo*.

### Evidence of lysosomal dysfunction in Slc4a11 KO corneal endothelial cells

Active lysosomes are essential for the ultimate degradation of autophagy substrates. With the observation of aberrant autophagy flux in KO MCEC, we investigated whether lysosomal function is normal in these cells. Western Blot analyses indicate decreased expression of two key lysosomal membrane proteins, Vacuolar ATPase (v-ATPase) and acid hydrolase Cathepsin B in KO MCEC (FIG 2A and B). We next determined whether the master transcription regulator of lysosomal function and biogenesis, Transcription factor EB (TFEB) ^18^, is affected by *Slc4a11* KO. TFEB levels were significantly decreased in KO MCEC compared to WT (FIG 2A and B). We also examined the levels of Co-ordinated Lysosomal Expression And Regulation (CLEAR) pathway genes ^19,20^. These genes regulate various stages of autophagosome generation, lysosomal biogenesis/function, and are transcriptionally regulated by TFEB. Figure 2C shows that CLEAR pathway genes were significantly downregulated in KO MCEC consistent with the lack of TFEB activity in these cells. Using immunofluorescence of Lysosome associated membrane protein 1 (Lamp1), we found that Lamp1 fluorescence intensity (arbitrary units) was significantly reduced in KO MCEC (75±23 au) when compared to WT MCEC (115±12 au) (FIG S2A and B). These results suggest that lysosomal numbers may be decreased in MCEC KO cells.

**Figure 2.**
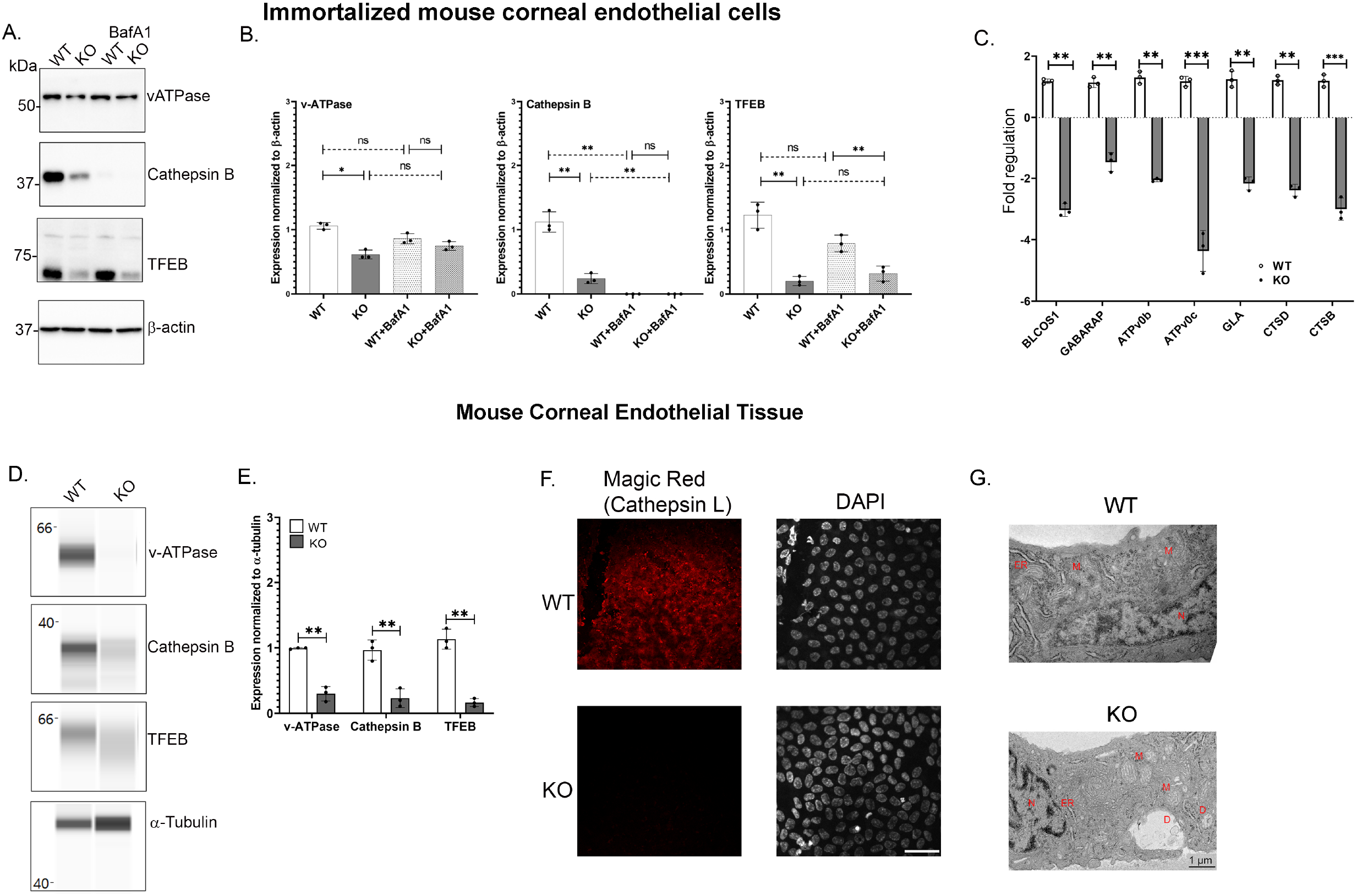
Lysosomal dysfunction is evident in Slc4a11 KO corneal endothelial cells. **Panel A** Western Blots of *Slc4a11* WT and KO Mouse Corneal Endothelial Cells (treated with or without 50nM BafA1) showing the expression of v-ATPase, Cathepsin B, and TFEB. **Panel A** Quantification of Western Blots from Panel A. n=3. *p<0.01, **p<0.001, ns-not significant **Panel A** Q-PCR showing the expression of a subset of TFEB regulated CLEAR network genes in *Slc4a11* WT and KO MCEC. BLOCS1 – Biogenesis of Lysosomal Organelles Complex1 Subunit1, GABARAP - GABAA receptor-associated protein 1, ATPv0b – vATPase subunit b, ATPv0c - vATPase subunit C, GLA –Alpha galactosidase A, CTSD – Cathepsin D, and CTSB-Cathepsin B. **Panel A** WES analysis of corneal endothelial tissues from *Slc4a11* WT and KO animals detecting the levels of v-ATPase, Cathepsin B, and TFEB. **Panel A** Quantification of WES runs from Panel D. n=3, **p<0.001 **Panel A** Magic Red staining to assess Cathepsin L activity in *Slc4a11* WT and KO corneal endothelial tissue. Scale bar – 50μm **Panel A** Electron microscopy images of *Slc4a11* WT and KO corneal endothelial tissue. D-Debris laden double membrane structures, ER-Endoplasmic reticulum, M-Mitochondria, and N-Nucleus.

Next, we examined lysosomal function in corneal endothelial tissue of *Slc4a11* KO animals. We observed a reduction in the expression of lysosomal proteins (FIG 2D and E), similar to that seen in MCEC. We then determined whether lysosomal Cathepsin L activity is affected in *Slc4a11* KO mice corneal endothelia. We used the Magic Red assay, which results in red fluorescence following Cathepsin digestion. Our data show decreased levels of Magic Red fluorescence in the KO tissue when compared to age matched WT controls (FIG 2F). Lastly, electron microscopy analyses of *Slc4a11* KO corneal endothelial tissue revealed the presence of debris laden structures, which were not observed in the WT tissues (FIG 2G). Lack of active lysosomes is consistent with the accumulation of undigested debris.

### ER stress and Unfolded Protein Response is activated in Slc4a11 KO corneal endothelia

The endoplasmic reticulum (ER) plays a crucial role in protein folding and lysosome production. Misfolded proteins trigger the unfolded protein response (UPR) through three main sensors, IRE1-α, PERK and ATF6. Activation of UPR leads to increased expression of ER chaperone protein, binding immunoglobulin protein (BIP), ER-related apoptosis protein, and growth arrest DNA damage protein (GADD) (FIG 3A) ^21^. Previous work has identified ER dysfunction in FCED ^22^, and we further investigated whether mitoROS in *Slc4a11* KO MCEC can affect ER stress.

**Figure 3.**
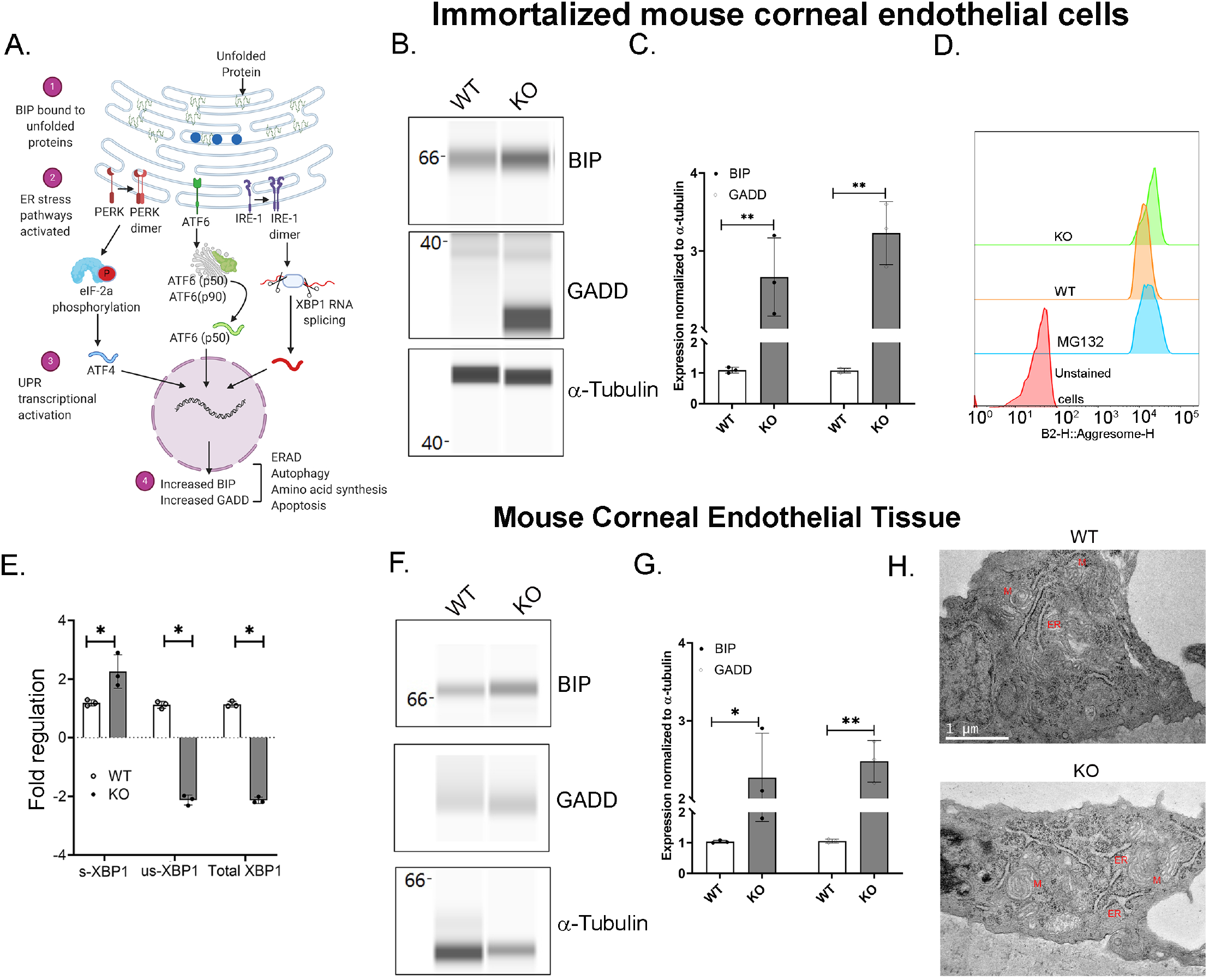
Presence of ER stress in Slc4a11 KO cells. **Panel A** ER stress through the unfolded protein response pathway is highly regulated through three main signaling pathways – ATF6, IRE1, and PERK. In a normal cell, the protein BIP is associated with the above mentioned molecules, thereby preventing the activation of ER stress associated signal transductions. With the accumulation of unfolded proteins, BIP is sequestered into these moieties (1), the stress pathways are activated (2), and ER stress associated transcriptional machinery is activated (3). Increased GADD and BIP levels are considered to be two of the major outcomes of ER stress pathways (4). **Panel A** WES analysis to determine the expression of ER stress markers, BIP and GADD in *Slc4a11* WT and KO MCEC. **Panel A** Quantification of Western Blots from Panel D. n=3, **p<0.001 **Panel A** Flow cytometry analysis of WT and KO MCEC to determine the presence of aggresomes (marker for unfolded proteins in the ER). APF (Aggresome Propensity Factor) was calculated to be 30.4 in cells treated with MG132, a proteostat inhibitor. APF was significantly higher in KO (11.4) compared to WT (1). N=3. **Panel A** Q-PCR results showing the transcript levels of spliced-XBP1 (s-XBP1), unspliced-XBP1 (us-XBP1) and total XBP1 levels in *Slc4a11* WT and KO MCEC. **Panel A** WES analysis to assess the levels of BIP and GADD proteins in *Slc4a11* WT and KO corneal endothelial tissues. **Panel A** Quantification of Western Blots from Panel D. n=3, **p<0.001 **Panel A** Electron micrographs of *Slc4a11* WT and KO corneal endothelial tissue. M-Mitochondria, ER – Endoplasmic Reticulum.

Increased expression of BIP and GADD, were observed in *Slc4a11* KO cells (FIG 3B and C). To determine the levels of unfolded proteins present in our samples, we conducted flow cytometry analysis of aggresomes, an accumulation of misfolded proteins following cellular stress ^23^, in MCEC WT and KO samples. Our analysis revealed significantly increased levels of aggresomes in the KO cells when compared to WT (FIG 3D).

Splicing of the gene, X-Box Binding Protein-1 (XBP1), occurs in response to the IRE-1α pathway. Using primers that can bind to unspliced XBP1, spliced XBP1 and Total XBP1 ^24^, we conducted real time PCR. Compared to WT samples, KO cells had increased levels of the spliced XBP1 transcripts (FIG 3E), whereas the unspliced and total XBP1 levels were decreased in KO MCEC.

In the mouse corneal endothelium, we observed increased BIP and GADD expression in 10 week old *Slc4a11*^-/-^ animals but not in the age matched *Slc4a11*^+/+^ animals (FIG 3F and G). Next, we conducted electron microscopy to determine whether there are any structural changes in the ER of the corneal endothelia of *Slc4a11* KO animals. Ten week old *Slc4a11* KO mice corneal endothelia reveal the presence of ER with dilated lumen that was not observed in WT samples of age matched animals (FIG 3H).

### Treatment with mitochondrial ROS quencher, MitoQ, reverses ER stress, autophagy impairment, and lysosomal dysfunction in Slc4a11 KO corneal endothelial cells

Studies from our lab and others have identified mitochondrial ROS to be the major cause of cell death in KO MCECs ^5,6^. In addition, mitochondrial ROS can also cause lysosomal dysfunction and ER stress in some systems ^25–28^. Therefore, we asked if quenching mitochondrial ROS will impact autophagy flux, lysosomal dysfunction and ER stress in *Slc4a11* KO corneal endothelial cells. We have previously shown that *Slc4a11* KO corneal endothelial cells or mouse corneal endothelial tissue have increased mitochondrial ROS ^5^ and this was confirmed for the current study (FIG S3A and B). Treatment with 2μM MitoQ, a mitochondrially directed ROS quencher, resulted in a significant reduction in mitochondrial ROS in the KO MCEC (FIG S3A and B).

Next, we determined whether autophagy induction was affected by the treatment with MitoQ. Wes analysis revealed reduced expression of p-mTOR in *Slc4a11* KO cells with MitoQ while the mTOR levels remained comparable in both WT and KO samples (FIG 4A and B). Moreover, with MitoQ treatment, P62 levels were significantly reduced in KO cells (FIG 4A and B). As a complementary approach to measure autophagy flux, we transfected WT and KO cells with pMRX-IP-GFP-LC3-RFP-LC3ΔG, followed by the treatment of MitoQ. We observed decreased GFP/RFP ratio in the WT cells, and elevated GFP/RFP levels in the KO cells (FIG 4C and D). With MitoQ treatment, KO cells showed a significantly reduced GFP/RFP levels that were comparable to those of WT cells (FIG 4C and D). These data strongly suggest the restoration of autophagy flux in KO MCEC with the quenching of mitochondrial ROS.

**Figure 4.**
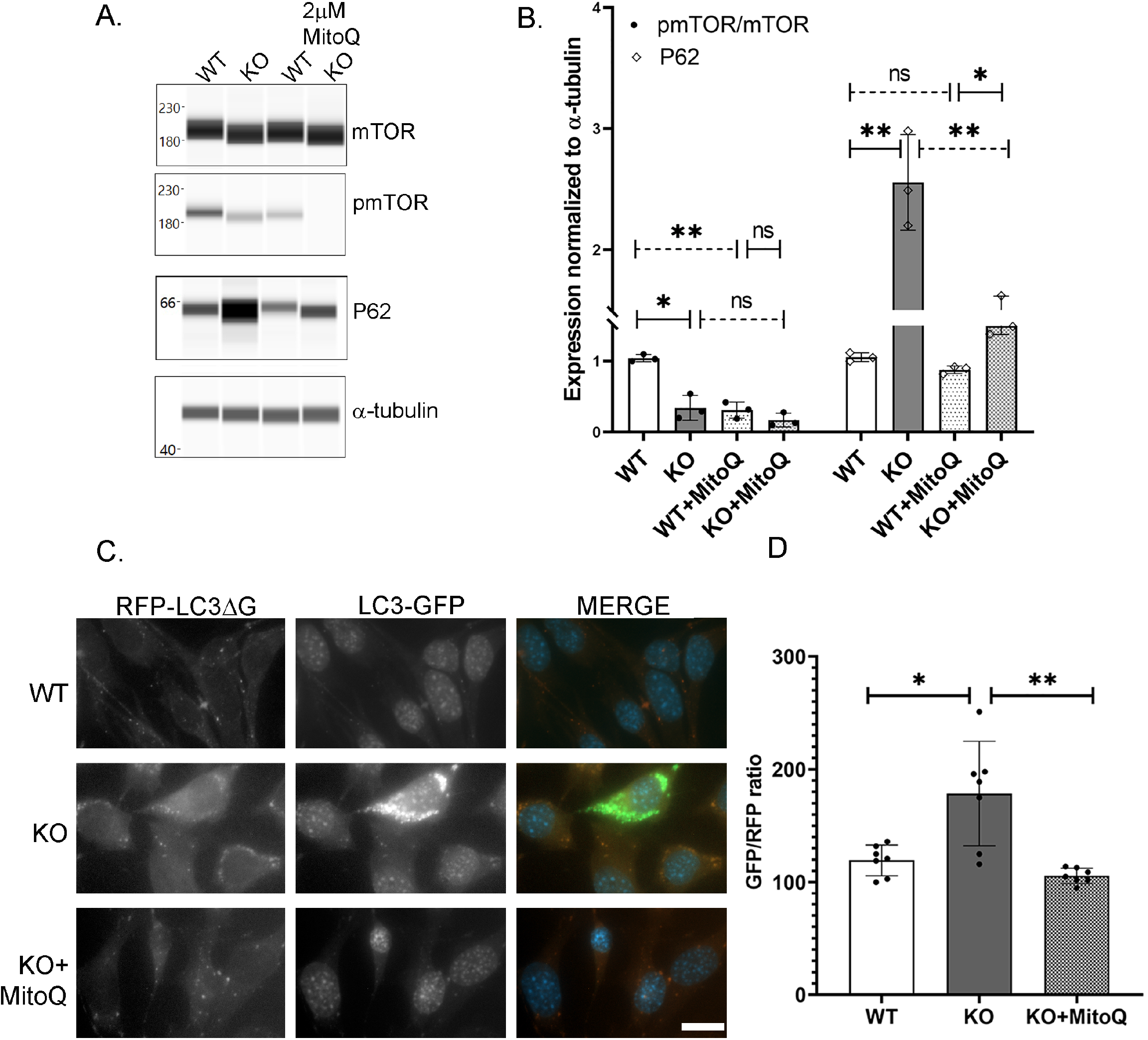
Treatment with MitoQ increases autophagy flux in MCEC KO cells. **Panel A** WES analyses to detect the expressions of pmTOR, mTOR and P62 in WT and KO *Slc4a11* MCEC WT and KO cells with MitoQ treatment. **Panel A** Quantification of Western results from Panel C. n=3, *p<0.01, **p<0.001, ns – not significant **Panels A** Detection of GFP and RFP fluorescence ratio in Slc4a11 WT, KO (+/-MitoQ) MCEC after transfection with GFP-LC3-RFP-LC3ΔG. Scale bar – 20 μm **Panel A** Quantification of fluorescence ratio from Panel C. *p<0.01, **p<0.001

The effect of MitoQ on lysosomal function was determined using Wes analysis of v-ATPase and TFEB. The expression levels of both these lysosomal genes were increased in KO MCEC with MitoQ treatment (FIG 5A and B). We also found that with MitoQ treatment, Lamp1 intensity was significantly increased in MCEC KO (FIG 5C and D). In addition, we also observed increased expression of CLEAR pathway genes in MitoQ treated KO cells, indicative of the restoration of TFEB activity (FIG S4A). All of these data indicate that lysosomal function recovers as a result of MitoQ treatment in MCEC KO.

**Figure 5.**
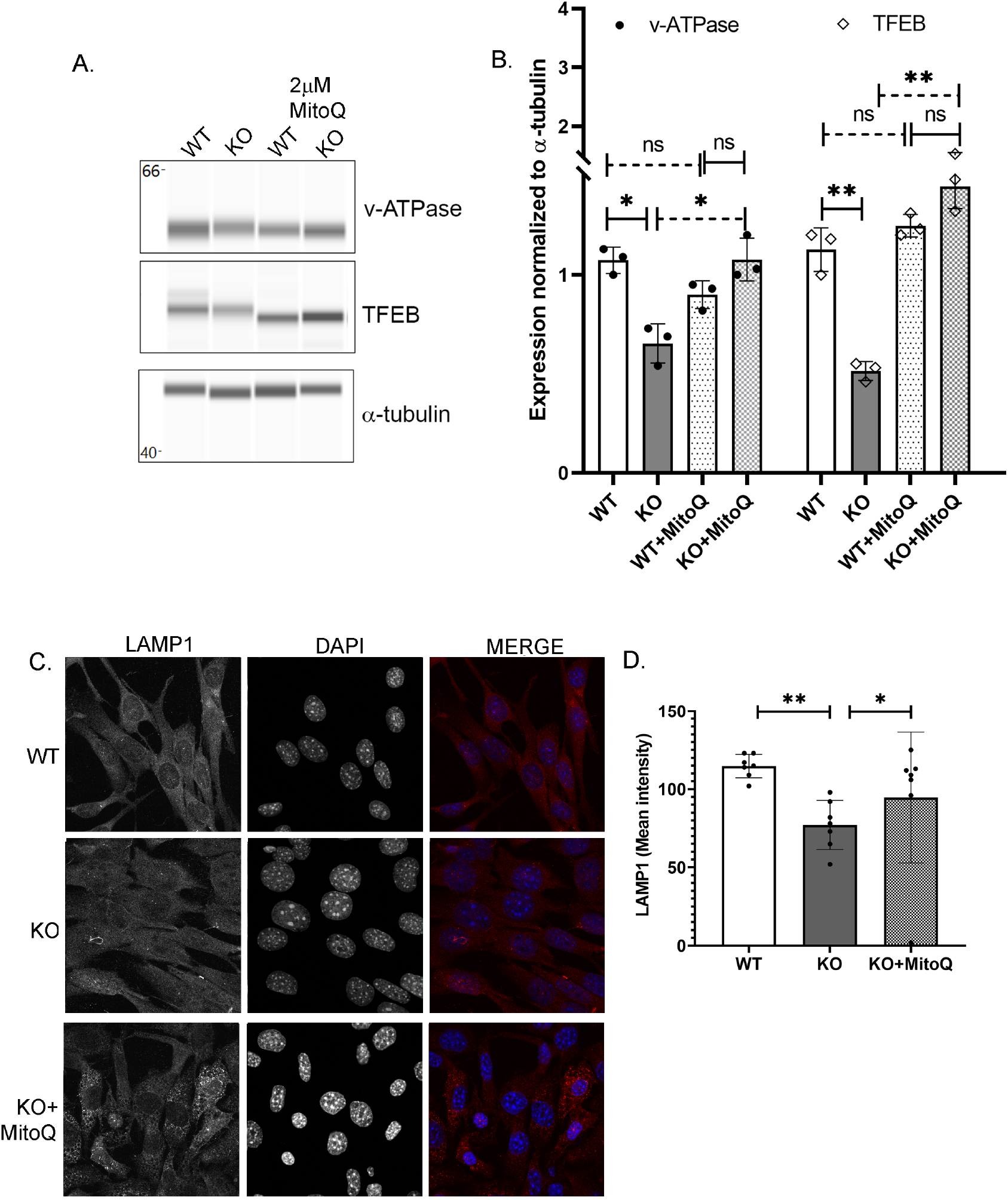
Treatment with MitoQ improves lysosomal function in MCEC KO cells. **Panel A** WES analysis results showing the expressions of expression of lysosomal proteins, v-ATPase, and TFEB in WT and KO *Slc4a11* MCEC. **Panel A** Quantification of Western Blot results from Panel C. n=3, *p<0.01, **p<0.001, ns – not significant **Panel A** Lamp1 staining in *Slc4a11* WT and KO (+/-MitoQ) MCEC. Scale bar - 5 μm **Panel A** Quantification of Lamp1 staining from Panel C. *p<0.01, **p<0.001

Next, we examined the effects of MitoQ on ER stress. Both BIP and GADD levels were significantly decreased in MCEC KO with the treatment (FIG 6A and B). Aggresome levels were analyzed by fluorescence microscopy using thapsigargin as a positive control. Without MitoQ treatment KO MCEC contained higher intensity of aggresomes (165±15), but in its presence these levels decreased significantly (105±12) (FIG 6C and D).

**Figure 6.**
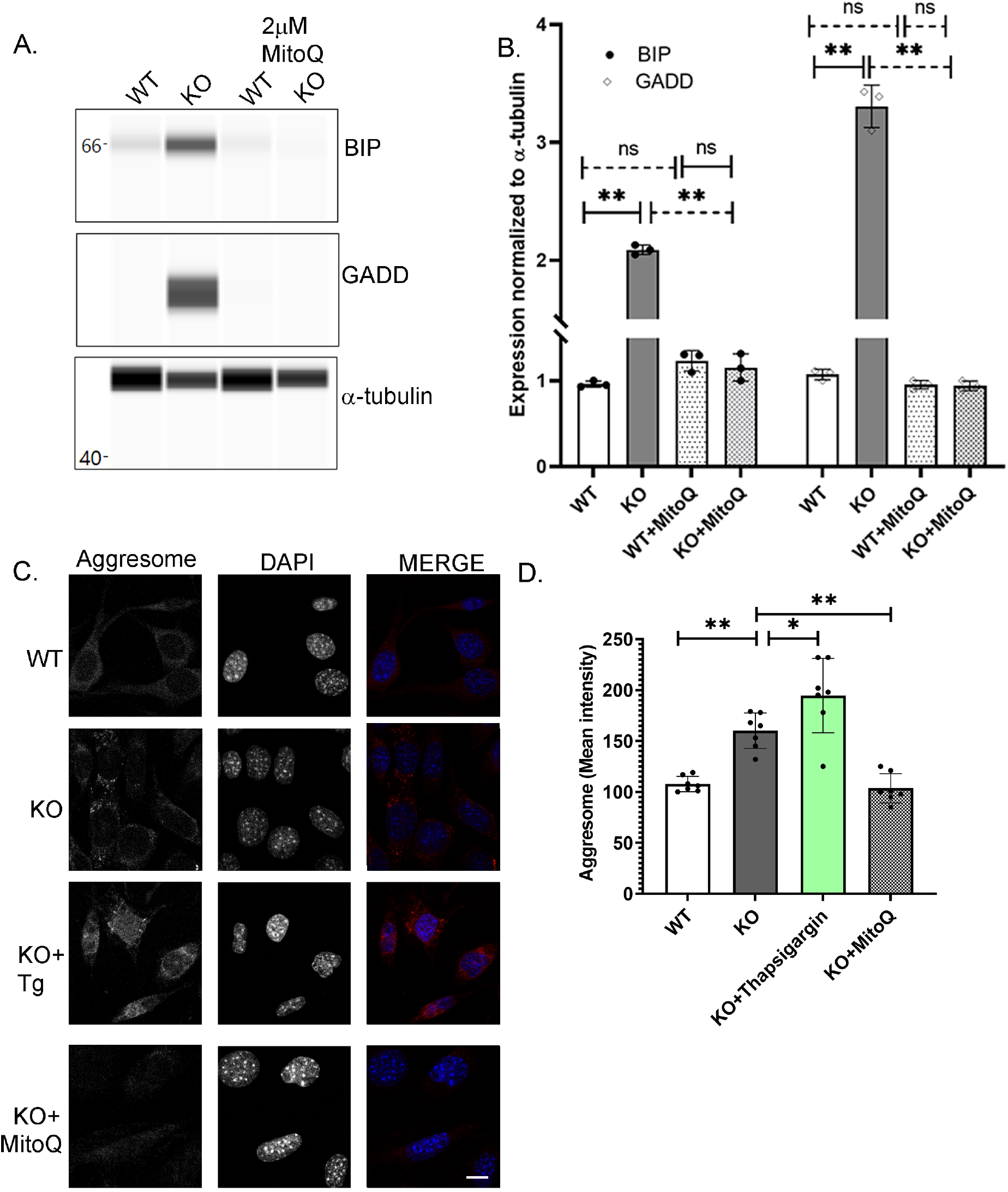
Treatment with MitoQ decreases ER stress in MCEC KO cells. **Panels A** WES analysis showing the expression of BIP and GADD in *Slc4a11* WT and KO Mouse Corneal Endothelial Cells. **Panel A** Quantification of Western results from Panel C. n=3, *p<0.01, **p<0.001, ns – not significant **Panel A** Aggresome staining in *Slc4a11* WT and KO (+/-MitoQ) MCEC. Cells were treated with Thapsigargin (0.1μM) as a positive control for aggresome staining. Scale bar - 5 μm **Panel A** Quantification of aggresome staining from Panel C. *p<0.01, **p<0.001

### MitoQ mediated reduction of Corneal Edema in Slc4a11 KO

The major clinical feature of CHED is progressive corneal edema, and the *Slc4a11* KO mouse model recapitulates this phenotype ^14^. In this animal model, significant corneal edema is apparent at eight weeks of age, and this continues to increase through adulthood. To determine if MitoQ could reverse the *in-vivo* corneal edema of *Slc4a11* KO mice, we conducted intra peritoneal injections of 2mM mitoQ on 6-week-old mice twice a week for two weeks. Our data show that injections resulted in significant reduction in corneal edema in Slc4a11 KO animals (FIG 7A and B). Corneal thickness continued to progress in vehicle injected animals and remained relatively unchanged in the WT animals (FIG 7A and B). In order to determine the specificity of MitoQ in reducing corneal edema, we stopped the injections for two weeks, and measured corneal thickness. In these animals, the corneal edema increased (FIG 7C).

**Figure 7.**
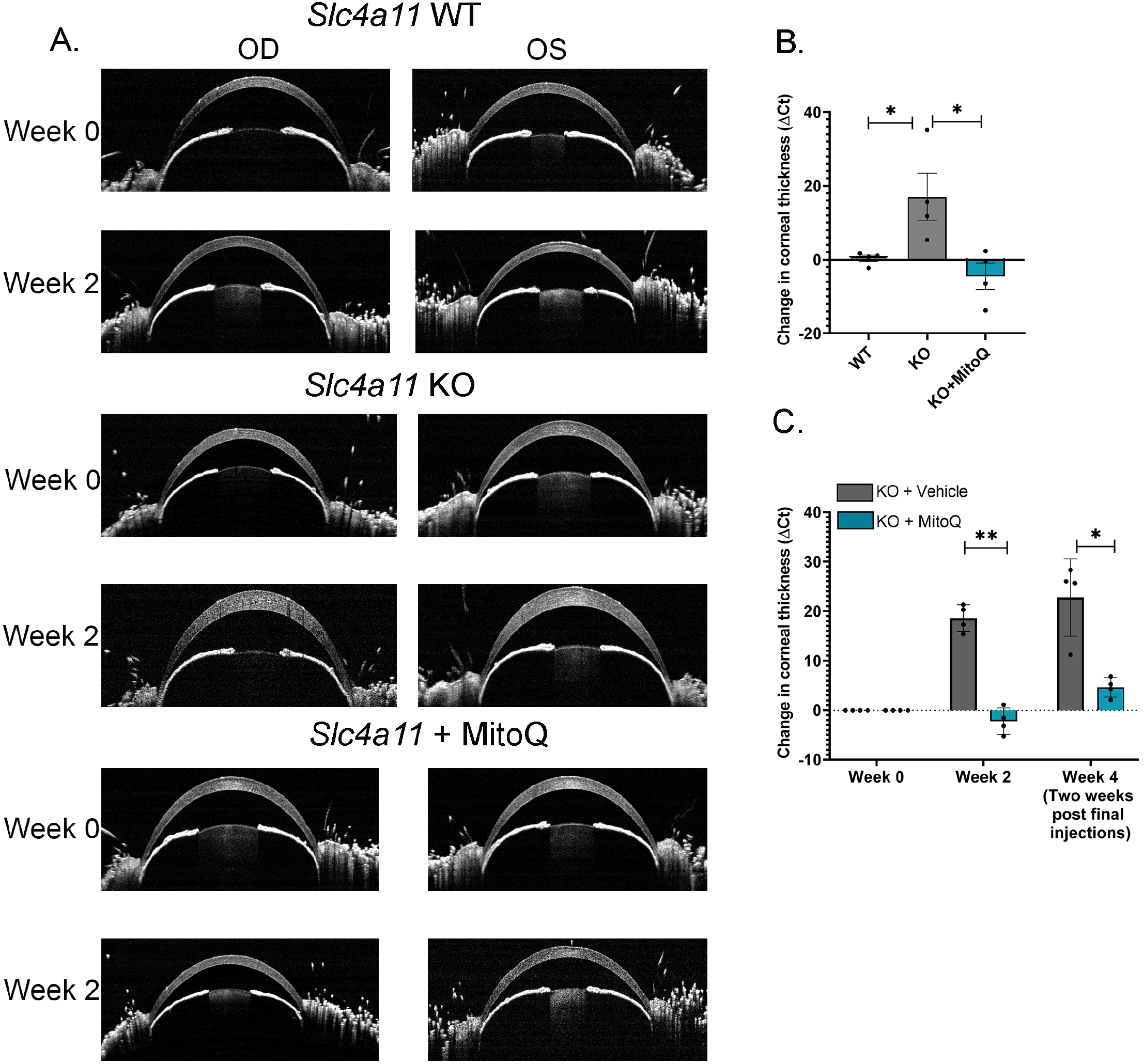
Intra-peritoneal injections of MitoQ partially rescues corneal edema in a mouse model of CHED. **Panel A** Optical coherence tomography (OCT) images of *Slc4a11* WT and KO mouse corneas. OCT measurements of corneal thickness were made before the start of injections (Week 0), and right after the two week duration of the injections (Week 2). **Panel A** Quantification of change in corneal thickness from Panel A. *p<0.01 **Panel A** *Slc4a11* KO animals were injected either with MitoQ or vehicle for two weeks. After this time, both sets of animals were left without any injections for another two weeks. KO animals injected with MitoQ showed decrease in corneal edema for the first two weeks but the edema returned once MitoQ treatment was stopped.

## Discussion

In the current study, we show that ROS produced as a result of the absence of mitochondrial Slc4a11 causes ER stress and a deficiency of lysosomal numbers and activity. Evidence for activated autophagy in MCEC *Slc4a11* KO was previously shown ^5^, however our in depth analysis here indicates that whereas autophagy is activated in KO cells, autophagy flux is inhibited due to lysosome deficiency. Parallel analysis from mouse corneal endothelial tissue confirm this conclusion. Furthermore, quenching mitochondrial ROS rescued the phenotype both *in vitro* and *in vivo*. These studies highlight the multiple organelle dysfunctions that can arise as a result of mitochondrial ROS in *Slc4a11* KO corneal endothelial cells.

Decreased ratio of p-mTOR/mTOR as well as increased levels of LC3II/I in *Slc4a11* KO corneal endothelia indicate that the autophagy initiation steps are present in these cells. This is consistent with previous studies that show defects in glutamine catabolism and decreased ATP production in KO ^5,29^, which would favor increasing autophagy via reducing p-mTOR/mTOR ratio. The accumulation of P62, however indicates a downstream impairment of autophagy flux, and the lack of significant change in P62 levels in the presence of Bafilomycin A1 are consistent with impaired autophagy flux. We provide evidence that the final step of autophagy, lysosomal digestion of phagophores, is dysfunctional.

TFEB controls lysosomal biogenesis ^19,30^. mTOR phosphorylation of TFEB inhibits translocation to the nucleus ^18,31^, so normally we would expect increased TFEB transcriptional activity in KO vs WT since the demand for autophagy is greater in KO. However, total TFEB and the TFEB activity (as evidenced by CLEAR pathway gene expression) are lower in KO cells, consistent with fewer lysosomes and poor lysosomal function. MitoQ partially rescues TFEB function with concomitant rescue of lysosomes. MitoQ also further decreases the p-mTOR/mTOR ratio in both WT and *Slc4a11* KO cells (Fig 4A), which would favor TFEB transcriptional activation. This has a small effect on TFEB in WT, where autophagy is already robust, but a significant recovery of TFEB expression in KO. These results suggest that the excess mitoROS in KO has detrimental effects on TFEB expression.

In addition to mTOR, TFEB is regulated through multiple pathways in several cell types. ERK 2 is capable of phosphorylating TFEB, and thereby preventing its nuclear translocation ^30^. Protein Kinase C and its isoforms, PKCα, PKCβ, and PKCδ affect lysosomal biogenesis through TFEB regulation ^32^. Alterations in this pathway can also lead to the accumulation of TFEB in the cytosol. Reduced TFEB activity could be caused by decreased overall expression, as seen in the Slc4a11 KO, and/or increased phosphorylation, which prevents nuclear translocation. Conversely, the phosphatase Calcinuerin targets TFEB ^33^, creating a complex yet well-regulated control of TFEB activity. Calcineurin contains iron and zinc in its active site, and these are sensitive to high ROS ^34^, suggesting that MitoQ not only boosts TFEB expression, but could facilitate nuclear translocation. Furthermore, in a Huntington’s disease model it was shown that PGC1-α, known for its regulation of mitochondrial biogenesis, can increase expression and activate TFEB by preventing oxidative stress ^35^. Whether one or more of these pathways are responsible for TFEB regulation in corneal endothelial cells will be explored in future studies.

While mitochondrial ROS can activate lysosomal biogenesis and increase autophagy flux in some cell types, ROS has also been shown to impede lysosomal acidification and autophagy in others ^36,37^. Acute mitochondrial stress has been attributed to increasing autophagy flux, whereas chronic stress can lead to lysosomal dysfunction and stall autophagy ^36^. In the corneal endothelial cells from *Slc4a11* KO animals, it is reasonable to hypothesize that the lysosomal dysfunction, autophagy impairment and ER stress are the outcomes of chronic stress. Whether autophagy and ER function are normal in the early stages of the disease and then progress to chronic mitochondrial stress and autophagy and ER dysfunction remains to be determined.

Apart from being the power house of the cell, mitochondria are considered to be a major signaling hub ^38^. Through signals, or by physical interaction, mitochondria relay and receive information from many organelles including the ER and lysosomes ^39^. ER is responsible for protein maturation and correct folding. ER and mitochondria are connected through Mitochondria associated ER membranes (MAMs) ^40^. The redox state within the cell affects ER protein folding machinery, and can result in Unfolded Protein Response (UPR) ^41^, a cascade of signaling events that arise during ER stress. Studies in FCED have detected ER stress and UPR ^22^. Here, we show that mitochondrial ROS, which could alter redox levels in ER, can also induce ER stress in the *Slc4a11* KO model of CHED.

Mitochondrial ROS is well-known as one of the major factors contributing to cell death in several corneal diseases such as keratoconus, FCED, CHED and granular corneal dystrophy ^42,43^. Several studies have identified mitochondrial dysfunction, or oxidative stress as the trigger behind cell death in each of these diseases ^3,4,44–46^. Our study further characterizes the cross-talk between mitochondria, lysosomes and ER in the corneal endothelial cells, and reveal that at least two important physiological processes (autophagy and ER function) are disabled by increased mitochondrial ROS. Furthermore, we show that normal mitochondrial function is essential for the maintenance of lysosome and ER function in the corneal endothelial cells. To the best of our knowledge, our study is the first to characterize inter-organelle relationships in cornea.

Recent clinical studies have shown the efficacy of MitoQ in improving vascular function in older individuals. Studies involving type-2 diabetes ^47^, kidney disease ^48,49^, and sensorineural hearing loss ^50^ have all shown promising results using MitoQ as a therapeutic agent. Our findings that MitoQ can ameliorate the progression of corneal edema in *Slc4a11* KO mice adds to this list. MitoQ was initially discovered for its use toward rescuing mitochondrial dysfunction in Parkinson’s disease ^51^, however clinical trials were ultimately unsuccessful. Since repurposing of old drugs can circumvent many years of additional research that may go into the development of a new drug, MitoQ stands as a promising candidate toward the treatment of diseases such as CHED.

## Conclusions

Slc4a11 is an ammonia sensitive proton transporter present in the inner mitochondrial membrane and in the plasma membrane. Loss of this protein function leads to blinding diseases such as CHED, and FCED. In the current study, we show that in the absence of Slc4a11, mitochondrial dysfunction causes abnormalities in other organelles, thereby compromising critical physiological processes such as autophagy and protein folding (FIG 8). We show that the downstream events that arise in the absence of Slc4a11 can be circumvented by the use of a mitochondrial specific ROS quencher, MitoQ, thereby providing a novel therapeutic target for treating corneal endothelial dystrophies.

**Figure 8.**
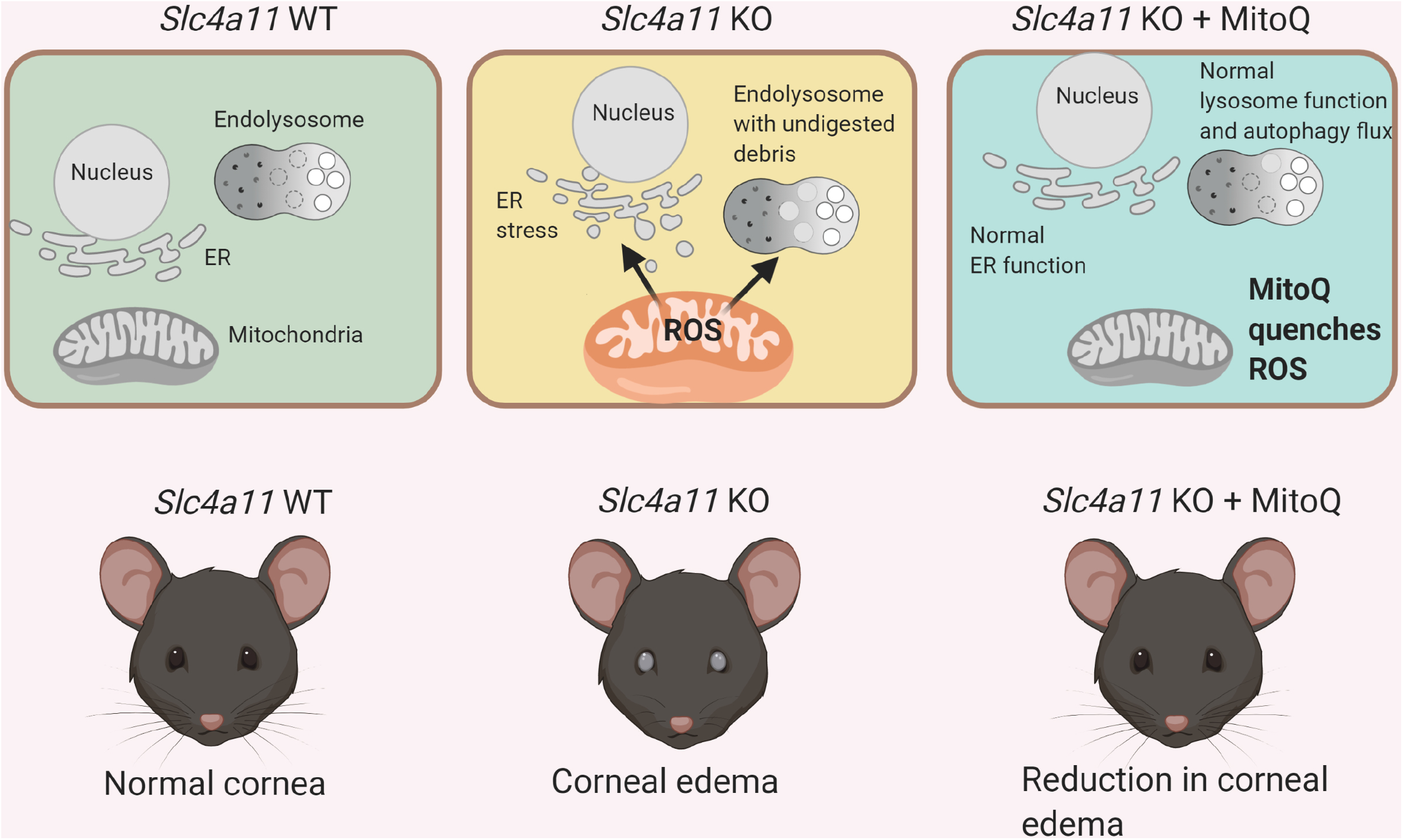
Graphical Abstract. In *Slc4a11* WT corneal endothelial cells, normal functions of mitochondria, ER and lysosomes were observed. However in the KO cells, mitochondrial dysfunction leads to increased ROS production. This in turn leads to ER stress, and lysosome dysfunction. With MitoQ treatment, *Slc4a11* KO cells show decreased ER stress, and normal lysosomal function. In Slc4a11 KO animals, corneal edema was partially rescued with MitoQ injections.

## Materials and Methods

### Animal model

*Slc4a11* KO mice were a gift from Eranga N. Vithana, Singapore Eye Research Institute ^14^. The mice were housed and maintained in pathogen-free conditions and used in the experiments in accordance with institutional guidelines and the current regulations of the National Institutes of Health, the United States Department of Health and Human Services, the United States Department of Agriculture and Association for Research in Vision and Ophthalmology (ARVO) Statement for the Use of Animals in Ophthalmic and Vision Research.

### Cell culture Experiments

The generation of conditionally immortalized mouse corneal endothelial cells (MCEC) Slc4a11^+/+^ and Slc4a11^-/-^ is described previously [12]. Cells were cultured in Complete Media, which contains OptiMEM-I medium (#51985; Thermo Fisher Scientific, Canoga Park, CA, USA), 14 mM Glucose and 4 mM L-AlanylGlutamine supplemented with 8% heat-inactivated fetal bovine serum (FBS) (#10082139; Thermo Fisher Scientific), EGF 5 ng/ml (#01-107 Millipore, Darmstadt, Germany), pituitary extract 100 μg/ml (Hyclone 15 Laboratories, Logan, UT, USA), calcium chloride 200 mg/L, 0.08% chondroitin sulfate (#G6737; SigmaAldrich Corp., St. Louis, MO, USA), gentamicin 50 μg/mL (#15710072; Thermo Fisher Scientific), antibiotic/antimycotic solution diluted 1:100 (#15240062; Thermo Fisher Scientific) and 44 units/mL IFN-γ (#485-MI; R&D Systems, Minneapolis, MN, USA).

For the experiments, cells were incubated in Assay Media, which contained Earle’s Balanced Salt Solution (no glutamine, no sodium pyruvate, no phenol red, contains 5.5mM glucose) (#141553; Thermo Fisher Scientific), supplemented with 0.5mM glutamine (#250030-081, Thermo Fisher Scientific) and 0.5% dialyzed FBS (#26400-036; Thermo Fisher Scientific) at 33°C for 16 hrs. Drug treatments, 50nM Bafilomyicn (#SML1661, SigmaAldrich Corp.), 2μM MitoQ (#317102, Medkoo Biosciences, Morrisville, NC, USA), or 0.1μM Thapsigargin (SML1845, Sigma Aldrich) were added into the assay media for 16 hours.

### Transient Transfection

MCEC WT and KO cells were grown in glass cover slips. Using Lipofectamine 3000 (L3000001, Thermo Fisher Scientific), following manufacturer’s instructions, cells were transfected with pMRX-IP-GFP-LC3-RFP-LC3ΔG (#84572, Addgene). 72 hours post transfection, cells were subject to treatment in assay media for 16 hours. Cells were fixed using 4% paraformaldehyde for 15 mins at room temperature, washed in PBS and mounted using prolong gold antifade mounting media (P36930, Thermo Fisher Scientific) and examined by Zeiss 15 Apotome microscope (Zeiss, Oberkochen, Germany).

### Western Blot analysis

Protein lysates were prepared using 1X RIPA analysis (10X RIPA buffer, #9806, Cell Signaling Technologies, Danvers, MA, USA) containing protease and phosphatase inhibitors (#5872S, Cell Signalling Technologies). Protein concentration was measured using BCA assay (#23227, Thermo Fisher Scientific), and 20 ug of proteins were resolved by either 8% SDS-PAGE or 15% for lower molecular weights, transferred on to Nitrocellulose membrane (#1620115, BioRad, Hercules, CA, USA), blocked using 5% milk in TBST for 1 hour followed by primary antibody incubation at 4°C for overnight using the following antibodies at the appropriate dilutions– LC3 (1:1000, #2775, Cell Signaling Technologies), Atg5 (1:1000, #12994, Cell Signaling Technologies), P62 (1:1000, # 23214, Cell Signaling Technologies), mTOR (1:1000, #2983S, Cell Signaling Technologies), pMTOR (1:1000, #5536S, Cell Signaling Technologies), v-ATPase (1:1000, #14617S, Cell Signaling Technologies), TFEB (1:1000, #A303673A, Bethyl, Montgomery, TX, USA), Cathepsin B (1:1000, #31718, Cell Signaling Technologies), β-actin (1:2000, #A5441, Thermo Fisher Scientific).

Protein lysates from corneal endothelial peelings were pooled from 4 animals, lysed in RIPA lysis buffer, and analyzed using the Protein Simple Wes System (Protein Simple, San Jose, CA, USA) following manufacturer’s instructions. The antibodies used only for the Wes analysis are as follows – GADD (#NB600-1353S, Novus Biologicals, Centennial, CO, USA), BIP (#C50B12, Cell Signaling Technologies), α-tubulin (#NB100-690, Novus).

### Real time PCR

Total RNA was isolated using RNA mini kit (#74104, Qiagen, Germantown, Maryland, USA). 1μg of RNA was used to prepare cDNA using a high capacity RNA to DNA kit (#4388950, Thermo Fisher Scientific). Previously published primer designs were used to amplify XBP1, us-XBP1, s-XBP1, and β-actin^24^. NCBI primer designing tool was used to design all other primers used for this study. Real time PCR was conducted using SYBR green dye using a BioRad CFX96 system. Relative quantitation was performed using 2^-ΔΔCt^ method against housekeeping gene. Fold Change (FC) is calculated as 2^-ΔΔCT^. The data is plotted as Fold regulation. Fold Regulation is equal to Fold change when Fold Change >1. When Fold Change <1, Fold Regulation = -1/Fold Change

### Electron microscopy

Samples were fixed with 2.5% glutaraldehyde (#16020, Electron Microscopy Sciences, Hatfield, PA, USA), 4% paraformaldehyde (#15710, Electron Microscopy Sciences), in 0.1 M sodium cacodylate buffer, pH 7.2 at 4° C and post-fixed with 1% osmium tetroxide (# 19150, Electron Microscopy Sciences) in 0.1 M sodium cacodylate buffer (#12300, Electron Microscopy Sciences), pH 7.2 at 4° C. Samples were dehydrated in a graded ethanol series to 100% ethanol, transitioned to propylene oxide (#20401, Electron Microscopy Sciences), and infiltrated with Embed 812 resin (#14120, Electron Microscopy Sciences). Infiltrated samples were placed in flat embedding molds and polymerized at 65 ° C for 18 hours. Resin blocks were cut with a diamond knife using a Leica Ultracut UCT ultramicrotome (Leica Systems, Buffalo Grove, IL). Sections were placed on 300 mesh copper TEM grids (#0300-CU, Electron Microscopy Sciences) and stained with saturated uranyl acetate in aqueous solution, and lead citrate. Stained sections were viewed with a JEM-1010 TEM (JEOL, Peabody, MA, USA) at 80 kV and photographed with a Gatan MegaScan 794 CCD camera or JEM-1400plus TEM (JEOL) with a Gatan OneView CMOS digital camera.

### Flow Cytometry

MCEC cultures in 12-well format were trypsinized, washed and fixed using 4% Paraformaldehyde. Following permeabilization, cells were stained using MitoSOX (#M36008, Thermo Fisher Scientific) or Aggresome detection kit (#ENZ51035, Enzo Life Sciences, NY) following manufacturer’s instructions. After filtration using 50 μm sterile CellTrics Filters (#04-004-2327, Sysmex, Gorlitz, Germany) flow cytometry analysis was conducted on MACSQuant VYB (Miltenyi Biotech, Germany) and 10,000 cells were collected per acquisition. For Aggresome detection, cells treated with proteasome inhibitor, MG132, served as a positive control. Data were analyzed with FlowJo software (FlowJo, LLC, Ashland, OR, USA)

### Immunofluorescence

Mouse corneas were isolated, and washed in ice cold PBS. Following fixation in 4% Paraformaldehyde for 10 mins, the tissues were permeabilized using 0.5% Triton-X100 for 10 min, blocked using 3% Donkey Serum for 1 hour, and incubated in primary antibody containing LC3b (#2775, Cell Signaling Technologies) for overnight at 4°C, followed by secondary antibody incubation for 1 hour at room temperature. The corneas were cut radially, and mounted for imaging.

For Lamp1 staining, the cells were fixed using 4% paraformaldehyde for 15 mins, permeabilized using 0.5% Triton X-100 for 10 mins, and blocked in 3% normal donkey serum for 1 hour. Primary antibody incubation using Lamp1 primary antibody (1:100, #NB120-19294, Novus) was conducted overnight at 4°C, followed by PBS wash, and secondary antibody incubation at room temperature for 1 hour. Cells were washed again in PBS and mounted using prolong gold antifade mounting media.

For aggresome staining, cells were fixed using 4% Paraformaldehyde for 10 mins. The staining was carried out as outlined in the Aggresome detection kit manual (#ENZ51035, Enzo Life Sciences).

Magic Red staining was conducted on fresh corneal cups using Magic Red Cathepsin L assay kit (#941, Immunochemistry Technologies, Bloomington, MN, USA) following manufacturer’s instructions. Images were acquired for LC3b and Lamp1 staining using Zeiss LSM 800 confocal microscope (Zeiss, Oberkochen, Germany). Magic Red staining images were acquired on Zeiss 15 Apotome microscope.

### Quantification of Confocal Fluorescence Images

Fluorescence microscopy images were analyzed and fluorescence quantified using ImageJ (U. S. National Institutes of Health, Bethesda, Maryland, USA). At least 35 cells were outlined for each condition, and cells were counted from 8 or more images. Experiments were repeated on three different samples, and background intensities were subtracted before calculating mean fluorescence intensities.

### Corneal thickness measurement

Corneal thickness was measured using iVue Optical Coherence Tomography (Optovue, Fremont, CA). All measurements were made at the same time of day to account for circadian rhythm based changes in corneal thickness. At least three measurements were made for each eye, and the average ± SD was used for all animals. All corneal thickness measurements were blinded.

### MitoQ injections

MitoQ (#317102, Medkoo Bisociences) was dissolved in equal amount of ethanol: distilled sterile water to obtain a final concentration of 2mM. 100 μL of the above solution was intraperitoneally injected into the animals for twice a week (Monday and Thursday) for two weeks at the same time of day. Control animals were injected with equal volumes of vehicle at the same time/day for the same duration. Animals were monitored for any changes in health or behavior during the experiment duration. All analyses of corneal thickness following the MitoQ injections were blinded.

### Statistical Analysis

All experiments were performed at least 3 times on different days. Error bars represent Mean ±SD. Statistical significance was calculated using unpaired t-tests. ANOVA was used to determine statistical significance if more than two groups were analyzed. Statistical analyses were conducted using Graph Pad Prism software (La Jolla, CA, USA).

### Model figures

The graphical abstract, and the model figures of autophagy, and ER stress were created with the aid of BioRender.com

## Author contributions

RS and JAB – Conceptualization, Design of experimental plan, Interpretation of results

RS – All experiments (apart from the ones listed below), data analysis and first draft of the manuscript

DGO – MitoSOX flow cytometry assay, Interpretation of results MC – Lamp1 staining

## Funding sources

JAB – R01EY031321 and R01 EY008834

RS – NIH/NCATS CTSI TL1 TR002531 and UL1 TR002529 (2018-2020) and Knights Templar Career Starter Grant (2019-2021)

## Acknowledgements

The authors are thankful to Shimin Li for plasmid purification of GFP-LC3-RFP-LC3ΔG and Edward Kim for excellent technical assistance. Barry Stein IUB Electron Microscopy facility; Christiane Hassel, Indiana University Flow Cytometry Core Facility; Prof. Mallika Valapala for helpful suggestions toward experimental design, interpretation of results, and critical reading of the manuscript; and Prof. Catherine Cheng for the use of Protein Simple WES machine, Zeiss LSM800 Confocal microscope and critical reading of the manuscript.

## Conflicts of Interest

RS – None, DGO – None, MC-None, JAB - None

## Figure Legends for Supplementary Figures

S1A-The mechanism of action of GFP-LC3-RFP-LC3ΔG. Once cleaved by Atg4, GFP-LC3 is degraded by lysosomal activity, whereas RFP-LC3ΔG remains in the cytoplasm. GFP/RFP ratio can be used to measure autophagy flux. High ratio indicates low autophagy flux and vice-versa. Model adapted from ^17^

S1B-WT and KO MCEC show the expression of GFP-LC3- RFP-LC3ΔG.

S1C-Quantification of GFP/RFP ratio.

S2A-Lamp1 staining in WT and KO MCEC cells. KO cells show reduction in LAMP1 expression compared to WT samples.

S2B-Quantification of LAMP1 mean intensity.

S3A-Flow cytometry results of MitoSox (marker for mitochondrial ROS) positive cells. WT and KO MCEC treated with 2μM MitoQ for 17 hours.

S3B-Quantification of flow cytometry results.

S4 – Real time PCR results showing CLEAR pathway gene expression in WT and KO MCEC treated with 2μM MitoQ for 17 hours. BLOCS1 – Biogenesis of Lysosomal Organelles Complex1 Subunit1, TFEB – Transcription Factor EB, CTSD-Cathepsin D, ATPV0b - vATPase subunit b

## Notes

### Competing Interest Statement

The authors have declared no competing interest.

