## Supplementary figures and images for "MitoROS due to loss of *Slc4a11* in corneal endothelial cells induces ER stress, lysosomal dysfunction and impairs autophagy"

### S1

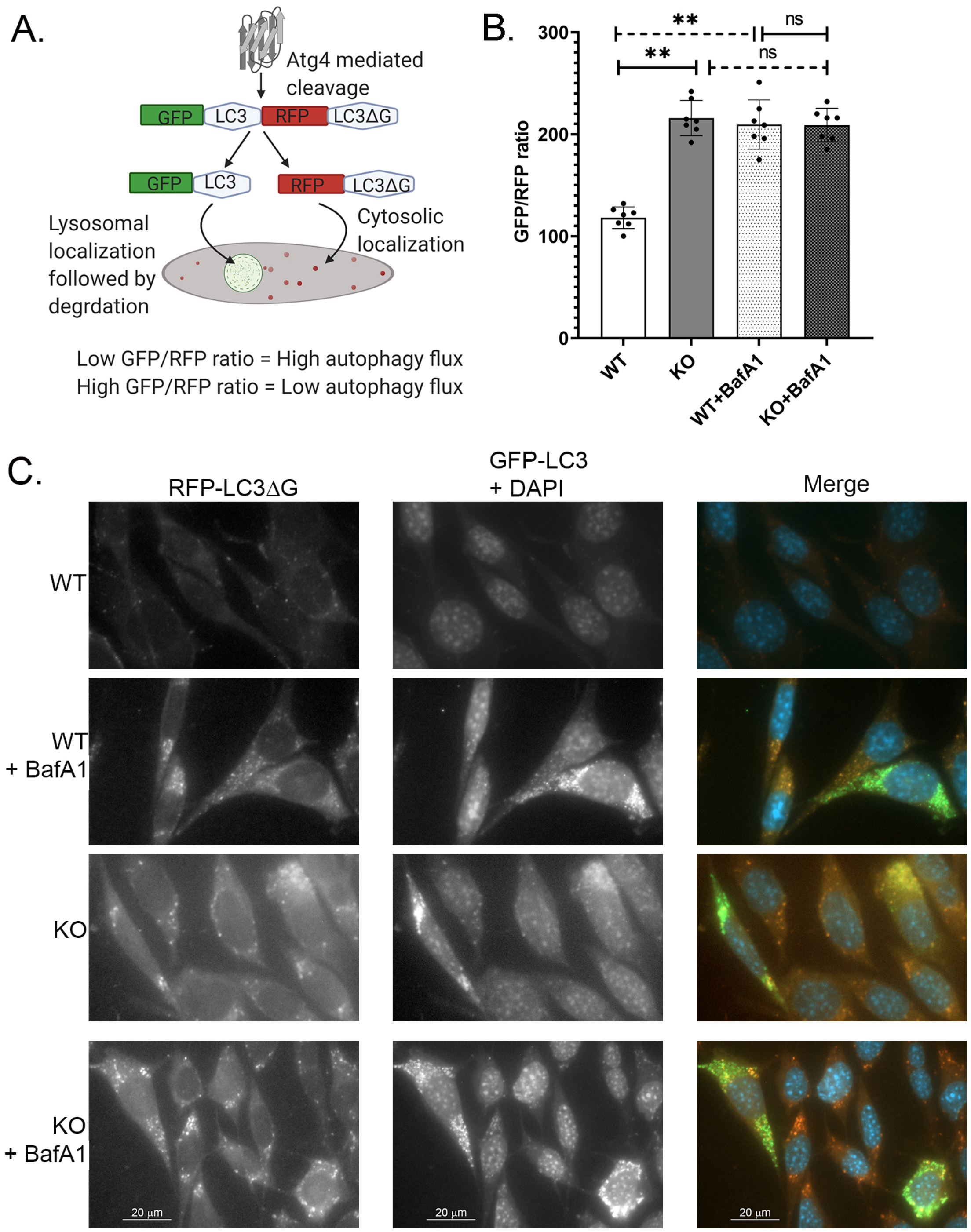

### S2

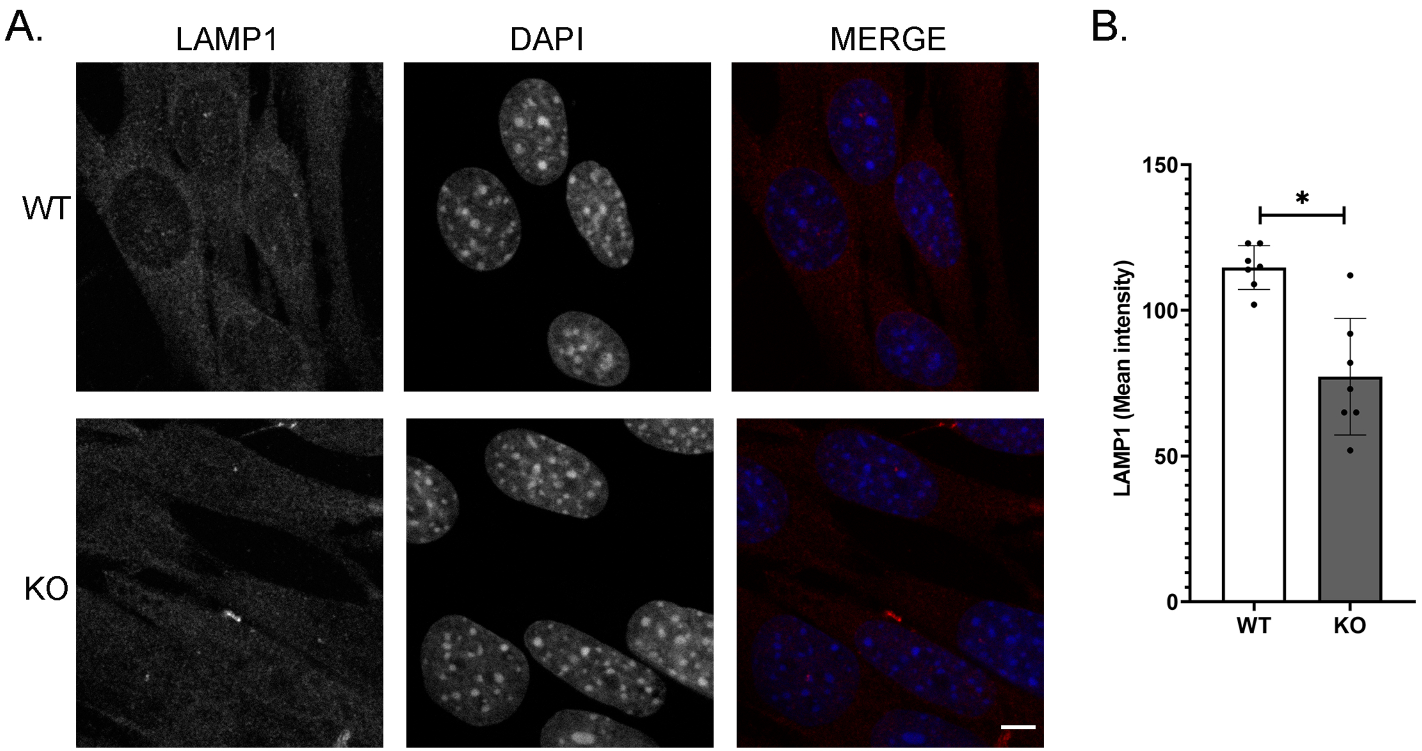

### S3

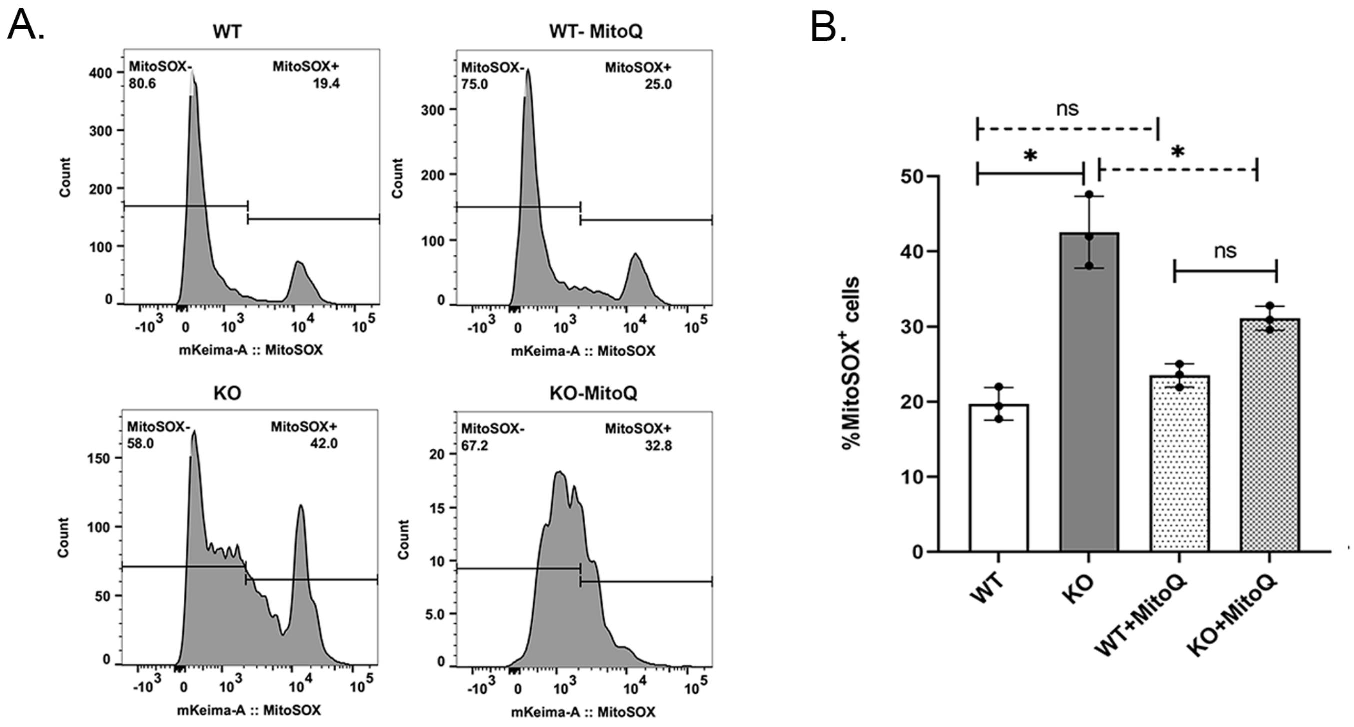

### S4

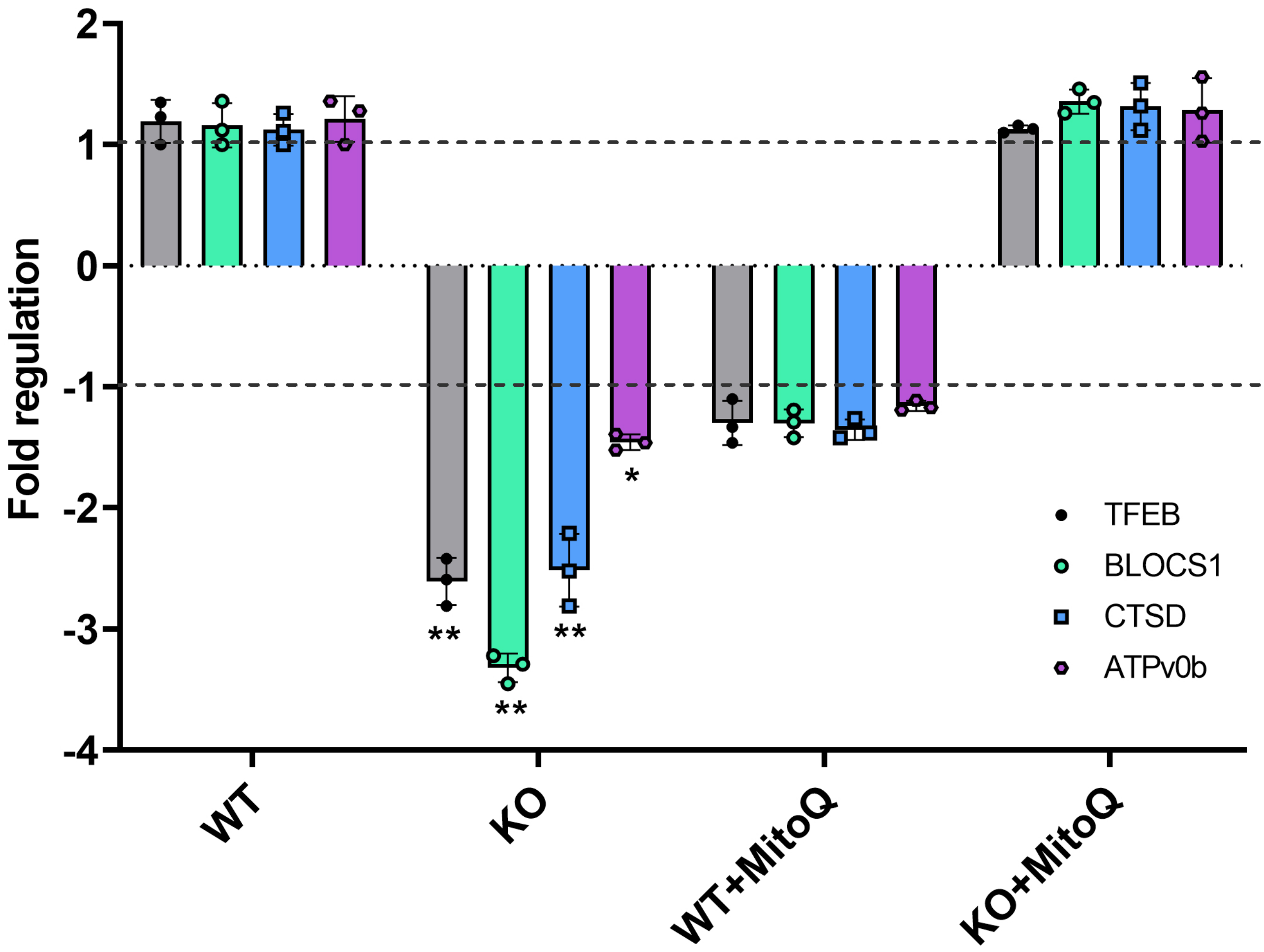
